# Determination of novel members in the *Drosophila melanogaster* anteriorposterior patterning system using natural variation

**DOI:** 10.1101/319434

**Authors:** Ashley A. Jermusyk, Sarah E. Gharavi, Aslesha S. Tingare, Gregory T. Reeves

## Abstract

The anterior-posterior axis of the developing *Drosophila melanogaster* embryo is patterned by a well-studied gene regulatory network called the Gap Gene Network. This network acts to buffer the developing pattern against noise, thereby minimizing errors in gene expression and preventing patterning defects.

In this paper, we sought to discover novel regulatory regions and transcription factors acting in a subset of the Gap network using a selection of wild-caught fly lines derived from the Drosophila Genetic Reference Panel (DGRP). The fly lines in the DGRP contain subtle genomic differences due to natural variation; we quantified the differences in positioning of gene expression borders of two anterior-poster patterning genes, *Krüppel* (*Kr*) and Even-skipped in 13 of the DGRP lines. The differences in the positions of *Krüppel* and Even-skipped were then correlated to specific single nucleotide polymorphisms and insertions/deletions within the select fly lines. Putative enhancers containing these genomic differences were validated for their ability to produce expression using reporter constructs and analyzed for possible transcription factor binding sites. The identified transcription factors were then perturbed and the resulting Eve and *Kr* positioning was determined. In this way, we found *medea, ultraspiracle, glial cells missing*, and *orthopedia* effect *Kr* and Eve positioning in subtle ways, while knock-down of *pangolin* produces significant shifts in *Kr* and subsequent Eve expression patterns. Most importantly this study points to the existence of many additional novel members that have subtle effects on this system and the degree of complexity that is present in patterning the developing embryo.

## Introduction

Spatial regulation of gene expression is of paramount importance in animal development, with improper regulation resulting in defects in development and disease states in adults [1,2]. Positional information is often initially delivered through a morphogen gradient [3,4]. Most generally, a morphogen is a molecule (usually a protein) that adopts a concentration gradient in space and that subsequently triggers expression of downstream patterning genes in a concentration-dependent fasion [3,4], typically through altering the activity of transcription factors in the affected cells’ nuclei. Further signaling between these downstream genes results in a web of interconnected genetic interactions called the genetic regulatory network (GRN) [5,6]. These networks are thought to buffer the developing pattern against noise, thereby minimizing errors in gene expression and preventing patterning defects [7–9]. Due to their importance in development, hours of laborious experimental work, computational methods, and genome-wide experimental methods such as ChIP-on-onchip have been invested to determine GRN topologies [10–14]. Even so, *it is thought that GRN maps remain incomplete* in even the best-characterized GRNs [8,15–18], suggesting that novel methods are required to discover unidentified components and DNA regulatory elements.

In this paper, we focus on the Gap GRN, which is responsible for patterning the anterior-posterior (AP) axis in the early *Drosophila melanogaster* embryo. This network is initiated by maternal factors, including Bicoid, Hunchback, Nanos, and Caudal [7,8,19]. *bicoid* (*bcd*) RNA is deposited by the mother at or near the anterior pole of the embryo and serves as a localized source of Bcd protein, which drives the establishment of an AP gradient of Bcd [20–22]. *nanos* (*nos*) RNA is deposited at the posterior pole of the embryo by the mother to create a Nos protein gradient opposite the Bcd gradient [23–25]. While both *caudal* (*cad*) and *hunchback* (hb) RNA are deposited ubiquitously by the mother, Bcd and Nos, respectively, act to create protein gradients [25,26]. These maternal inputs subsequently activate zygotic expression of the Gap genes - including *Krüppel* (*Kr*)*, knirps* (*kni*)*, hunchback* (*hb*), and *giant* (*gt*) -- which are expressed in broad stripes along the anteriorposterior axis [7,8,17]. Cross-repression between the gap genes serve to refine their borders [8,17,19]. The gap genes then activate the downstream pair-rule genes, which form the parasegments of the embryo [27] and control the expression of the segment polarity genes, which form the segments of the embryo [9,28].

Many of the currently-known connections within this network have been found via overt perturbations; however, we sought to find new connections within this network by quantifying subtle natural variation among wild-caught, in-bred lines that belong to the *Drosophila* Genetic Reference Panel (DGRP) [7,8,17,19,29]. Previous work has used the full DGRP panel, which consists of > 150 fly lines, to identify novel genes responsible for phenotypic changes in *Drosophila melanogaster* [29,30]. Three lines have been used to quantify variation in gene expression in AP patterning genes [31]. In this work, we focused on the subtle, but quantifiable natural variation in gene expression patterns of *Kr* and Even-skipped (Eve) in thirteen fly lines in the DGRP. Variations in gene expression domains were then linked to specific genomic differences between the lines. This study found how small genomic changes, even single nucleotide changes, outside of previously characterized enhancer regions can measurably impact gene expression patterns. We then used a position weight matrix approach, combined with literature ChIP-seq data, to identify possible transcription factor binding sites within these sites of genomic variation. We measured the expression domains of *Kr* and Eve in fly lines in which the identified transcription factors were perturbed, and found that *Kr* and Eve expression is altered subtly in four cases (*medea, ultraspiracle, glial cells missing*, and *orthopedia*) and overtly in a fifth case (*pangolin*). This evidence points to a larger number of unexplored genes that act within the early embryo to control anterior-posterior patterning.

## Results

### Measureable variation exists in Kr and Eve expression among wild-caught lines

The expression patterns of *Kr* mRNA and Eve protein were determined in 13 of the DGRP fly lines and in a laboratory control strain (yw; see Materials and Methods). *Kr* is expressed in a broad stripe 43 - 53% embryo length (Fig. 1A,C). Eve is expressed in seven narrow stripes (at 32, 40, 47, 54, 61, 68, and 77% embryo length) (Fig. 1B,D). We measured the expression patterns for both *Kr* and Eve (see Materials and Methods) in the mid-saggittal plane of the embryo. Variability among the lines was found in the positioning of the two *Kr* borders and Eve stripes 1 - 6 (ANOVA, p < 0.012) (Table 1, S1 Fig.). Due to the comparatively high variability within each line in positioning of Eve stripe 7, no statistically significant difference was found among the lines for that stripe (ANOVA, p = 0.08).

**Figure 1:**
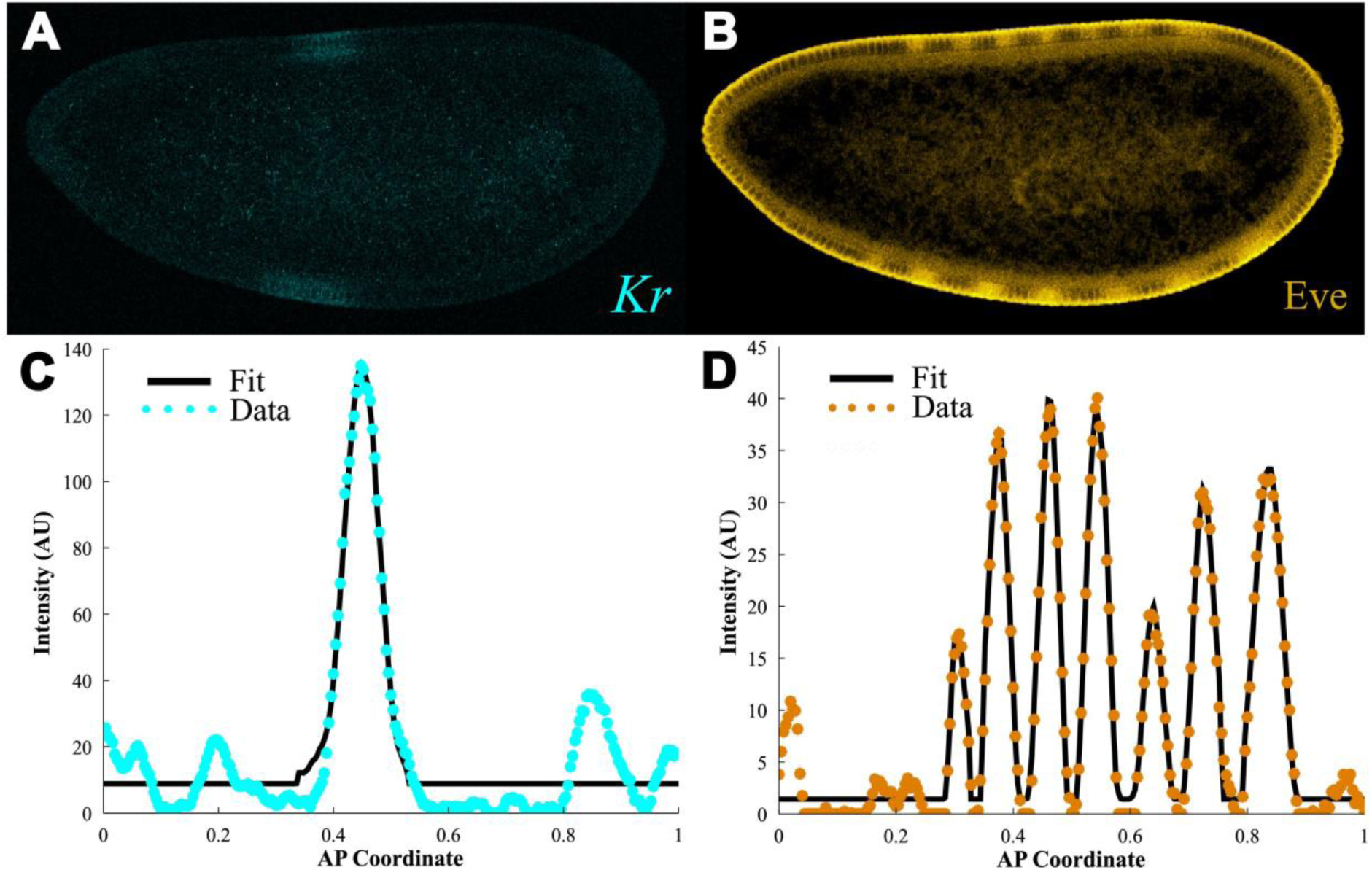
*Kr* and Eve expression quantified in the embryo reveals changes in gene expression among the fly lines. Normal expression of (A) *Kr* and (B) Eve as measured via *in situ* hybridization at the mid-saggittal section in an embryo (for these images and all images, anterior is to the left). Quantification of this expression along the dorsal half of this embryo where (*) is the normalized expression at each point and the solid black curve is the fit for (C) *Kr* and (D) Eve.

**Table 1:** ANOVA analysis of positioning of *Kr* and Eve across fly lines

| Border or Stripe | P-value (no control) | P-value (w/control) |
| --- | --- | --- |
| Kr (Anterior) | 2.36E-05 | 1.50E-06 |
| Kr (Posterior) | 4.28E-06 | 6.77E-07 |
| Eve Stripe 1 | 1.56E-06 | 6.29E-06 |
| Eve Stripe 2 | 1.98E-11 | 2.74E-11 |
| Eve Stripe 3 | 4.69E-11 | 1.09E-12 |
| Eve Stripe 4 | 1.54E-06 | 3.48E-12 |
| Eve Stripe 5 | 0.012 | 7.91E-13 |
| Eve Stripe 6 | 0.001 | 2.86E-05 |
| Eve Stripe 7 | 0.078 | 0.039 |

### Association Mapping to Locate Significant SNPs

Association mapping was used to determine if these differences in gene expression were correlated to specific genomic differences among the lines (Fig. 2A-B). All single nucleotide polymorphisms (SNPs) and short insertions and deletions (indels) 20 kb upstream and downstream of *Kr* and *eve* were evaluated; this region includes 661 SNP/indels near *Kr* and 646 near eve. A SNP or indel is taken as significant where there is a difference (twosided student’s t-test, p-value < 0.05; see Materials and Methods) between the position in the lines with the reference allele compared to the lines with the alternate allele (see S2 Fig.).

**Figure 2:**
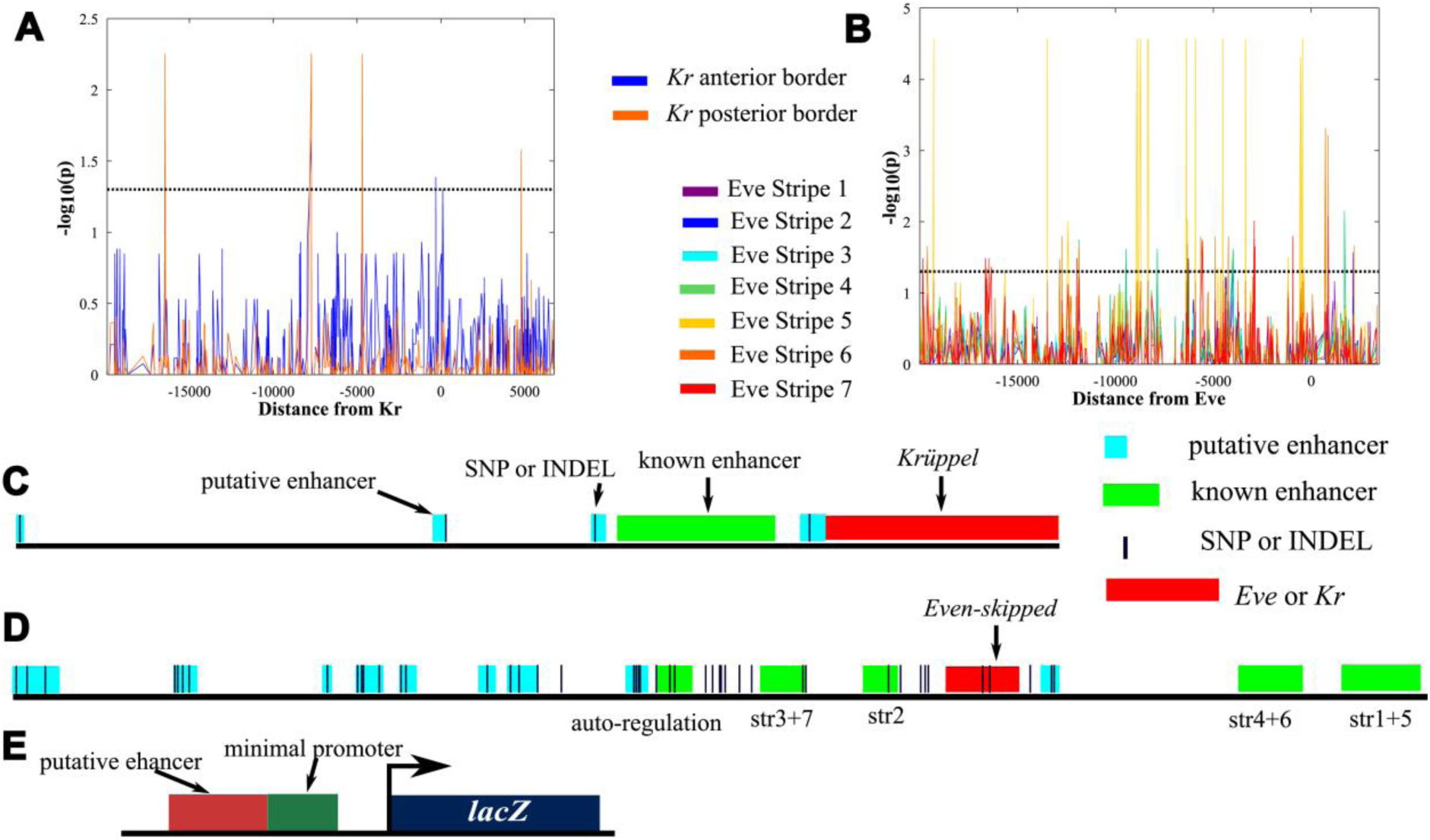
Results of association mapping analysis. Probability a given SNP or indel is correlated with changes in gene positioning for (A) *Kr* and (B) Eve. (C) Region surrounding the *Kr* gene with the SNPs and indels (thin dark blue bars) found to be associated with changes in *Kr* expression and the putative enhancers they were tested in. Known enhancers [34] are shown in green and putative enhancers tested are shown in cyan. (D) Locations of SNPs and indels found to be associated with changes in *eve* expression. Putative enhancer regions tested (cyan) and known enhancer regions (green, with the stripe regulated shown below, [10,27,32]) are shown. Both (C) and (D) are drawn to scale. (E) The putative ehancer plasmids contain the putative enhancer upstream of a minimal promoter. The enhancer, when active, drives expression of *lacZ*.

We found 5 statistically significant variants near the *Kr* locus, and 47 near eve. Of those near the *eve* locus, 13 were in known enhancers or within the *eve* coding sequence; since our desire was to screen for novel regulatory elements, these were not explored [10,27,32]. To screen for false positives and validate our findings, genomic regions between 161 and 1100 bp in size surrounding the statistically significant variants were tested for their ability to drive RNA expression *in vivo* using a reporter construct. These “putative enhancer regions,” which each contained one or more significant SNP/indels, were placed upstream of an *eve* minimal promoter to drive the expression of *lacZ* (Fig. 2) [33]. Four putative enhancers for *Kr* and twelve for *eve* were tested (where the “wild-type” allele for each variant was used - see Materials and Methods). The genomic positions of these regions and the primers used to create constructs with these regions are listed in S1 Table.

### Testing of Putative Enhancers

Of the 16 tested putative enhancers, three for *eve* and one for *Kr* were able to drive distinct expression *in vivo* (Fig. 3); representative embryo images are shown for all enhancers in S4 Fig. The minimal promoter used with these putative enhancers drives expression of a non-specific stripe at roughly 20–30% embryo length (Fig. 3I); only putative enhancers that generate expression outside of this region were explored further [33]. These expression patterns are relatively weak (Fig 3A, C, E, and G), which is to be expected since regulatory elements that drive overt gene expression patterns have already been identified by other methods.

**Figure 3:**
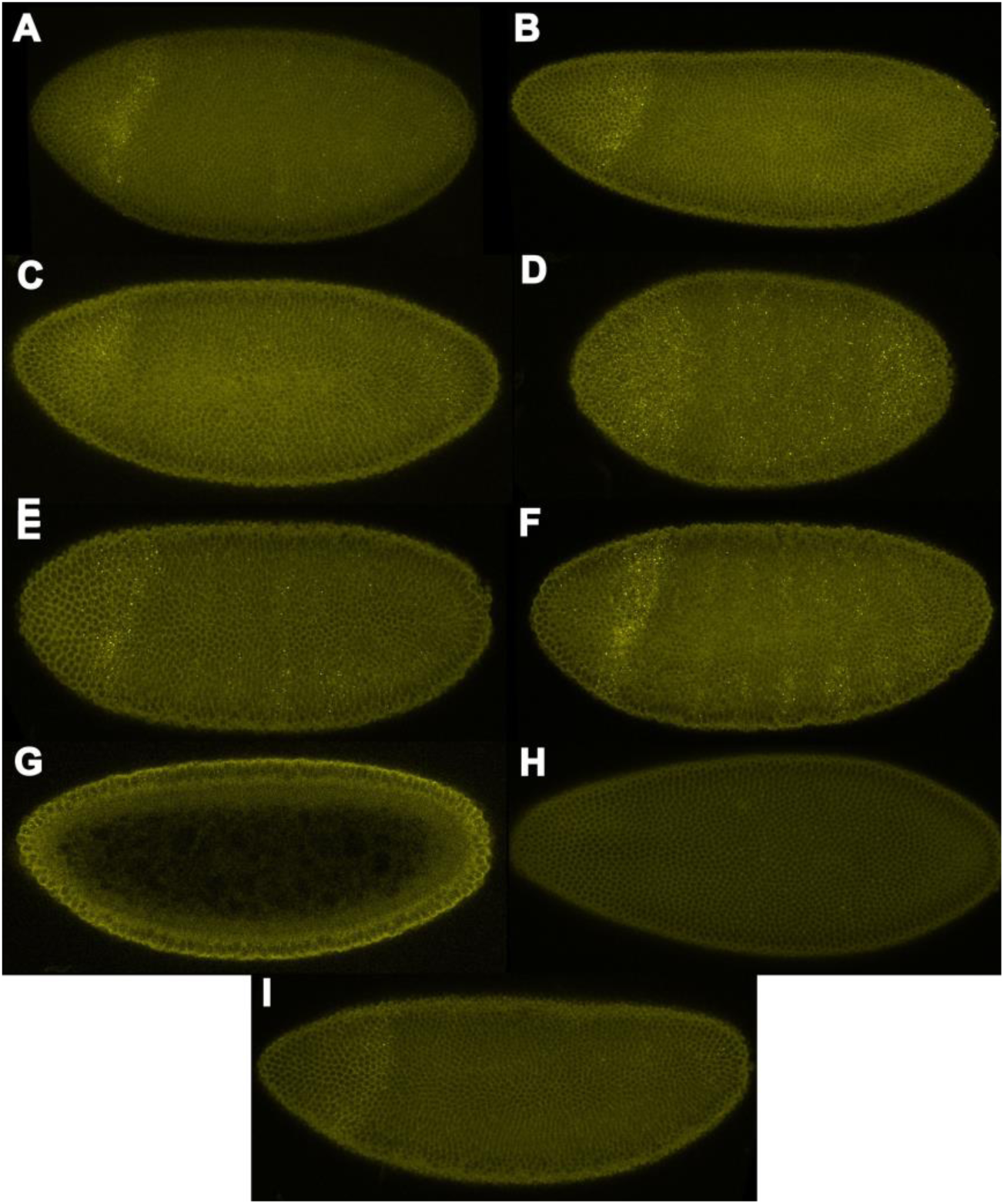
Results of Putative Enhancer Testing. (A) Expression of *lacZ* due to EveA enhancer is localized to the posterior region of the embryo. (B) In the mutated EveA enhancer expression is lost in this posterior region. (C) The EveB enhancer causes expression along the anterior and posterior poles of the embryo. (D) This anterior expression is not lost in the mutated EveB enhancer, in fact expression increases throughout the embryo. (E) The anterior pole and weak stripes of *lacZ* expression are driven by the EveC enhancer. (F) The expression in the stripes is increased in the mutated EveC enhancer, however expression in the anterior cap is lost. (G) The KrA enhancer drives expression at the anterior pole. (H) Expression is lost in the mutated KrA enhancer. (I) The putative reporter construct plasmid without any enhancer region (just the minimal promoter) drives a broad stripe of expression between 20–30% embryo length.

The putative enhancers that were able to produce expression *in vivo* in the context of the reporter constructs were mutated at the site of the SNP/indel to the alternate allele of this SNP/indel. These mutated enhancers produced altered expression patterns of the reporter gene (Fig. 3B,D,F, and H). For the EveA enhancer, the mutated version features an AACA deletion, which results in loss of the posterior expression found with the non-mutated enhancer. The change at the SNP within the EveB enhancer (from A to T) results in an increase in expression throughout the embryo. A loss in expression at the anterior pole of the embryo and simultaneously an increase in expression of stripes results from altering the EveC enhancer at the SNP (C to T). Mutating the indel (C insertion) in the KrA enhancer, results in a loss of expression in the anterior cap of the embryo.

### Determining novel transcription factors

This ability of these enhancers to drive expression and the change in these expression patterns when the SNP/indel is mutated may point to the presence of transcription factor binding sites within these putative enhancers at the SNP/indel. Therefore these enhancer regions were then analyzed using Position Weight Matrices to compile a list of transcription factor binding sites that may be present at the SNPs and indels of interest (S3 Fig., Table 2) [35–40,40–42]. Where available, ChIP data were also used to rule out or suggest transcription factors [35]. We ruled out for further investigation within this study any transcription factors already known to interact with the AP patterning system (see Table 2). The remaining transcription factors -- *glial cells missing* (*gcm*)*, medea* (*med*)*, orthopedia* (*otp*)*, ultraspiracle* (*usp*), and *pangolin* (*pan*) -- represent possible novel components of the AP network. We then tested these transcription factors for their ability to affect *Krüppel* and Even-skipped expression in mutant fly lines (see Materials and Methods) as compared to *yw* control expression patterns.

**Table 2:** Transcription factors identified as being likely candidates for binding to SNPs and indels of interest. Genes identified in bold were tested in this study.

| **Transcription Factor** | **Reporter construct** | **Suggested By** | **Notes** |
| --- | --- | --- | --- |
| Bcd | EveB | PWM | AP patterning gene [8] |
| Cad | EveA, EveB | ChIP | already known to interact [43] |
| D | EveA, EveB | ChIP | already known to interact [44] |
| Dl | EveA, EveC | ChIP | already known to interact [14] |
| En | EveB | PWM | AP patterning gene [45] |
| **Gcm** | **EveA** | **PWM** | **tested here** |
| Gt | EveB | PWM | AP patterning gene [8] |
| Hkb | KrA | PWM | eliminated by ChIP data |
| Kni | KrA | PWM | AP patterning gene [8] |
| Kr | KrA | PWM | AP patterning gene [8] |
| **Med** | **EveA, EveB** | **ChIP** | **tested here** |
| **Otp** | **KrA** | **PWM** | **tested here** |
| **Pan** | **EveB** | **PWM** | **tested here** |
| Slp1 | EveA, EveB | ChIP | already tested [46] |
| Sna | EveA | PWM | eliminated by ChIP data |
| Tin | KrA, EveA | PWM | effects eve [47] |
| Tll | EveC | PWM | eliminated by ChIP data |
| Twi | EveA, EveB | ChIP | already tested [7] |
| **Usp** | **EveC** | **PWM** | **tested here** |

### gcm effects positioning of Kr and Eve stripes 6 and 7

Enhancer EveA, which drives allele-specific expression in the posterior of the embryo, contains a possible transcription factor binding site for Glial cells missing (Gcm) at the site of the indel (per PWM analysis). The positioning of *Kr* and Eve stripes 6 and 7 (found in the posterior of the embryo) was found to be effected by knock-down of *gcm* (*glial cells missing*). We tested three different *gcm* RNAi lines, and found posterior shifts in the *Kr* domain and in the Eve 6 and 7 domains (see Fig. 4, S3 Table, S4 Table, and Materials and Methods). This shift in the positioning of Eve stripe 6 is consistent with the shift in Eve stripe 6 correlated with the indel in the EveA enhancer. However, *gcm* is known to be expressed in the anterior half of the embryo (15–35% embryo length on the ventral side) [48]. This would normally suggest an indirect effect on *Kr* and *eve* by Gcm through some other intermediary signal; however, our data suggest that Gcm directly interacts with *eve* and *Kr*. Therefore, it is possible that Gcm diffusion, combined with a Gcm binding partner expressed towards the posterior of the embryo, could be responsible for the effect on *Kr* and Eve.

**Figure 4:**
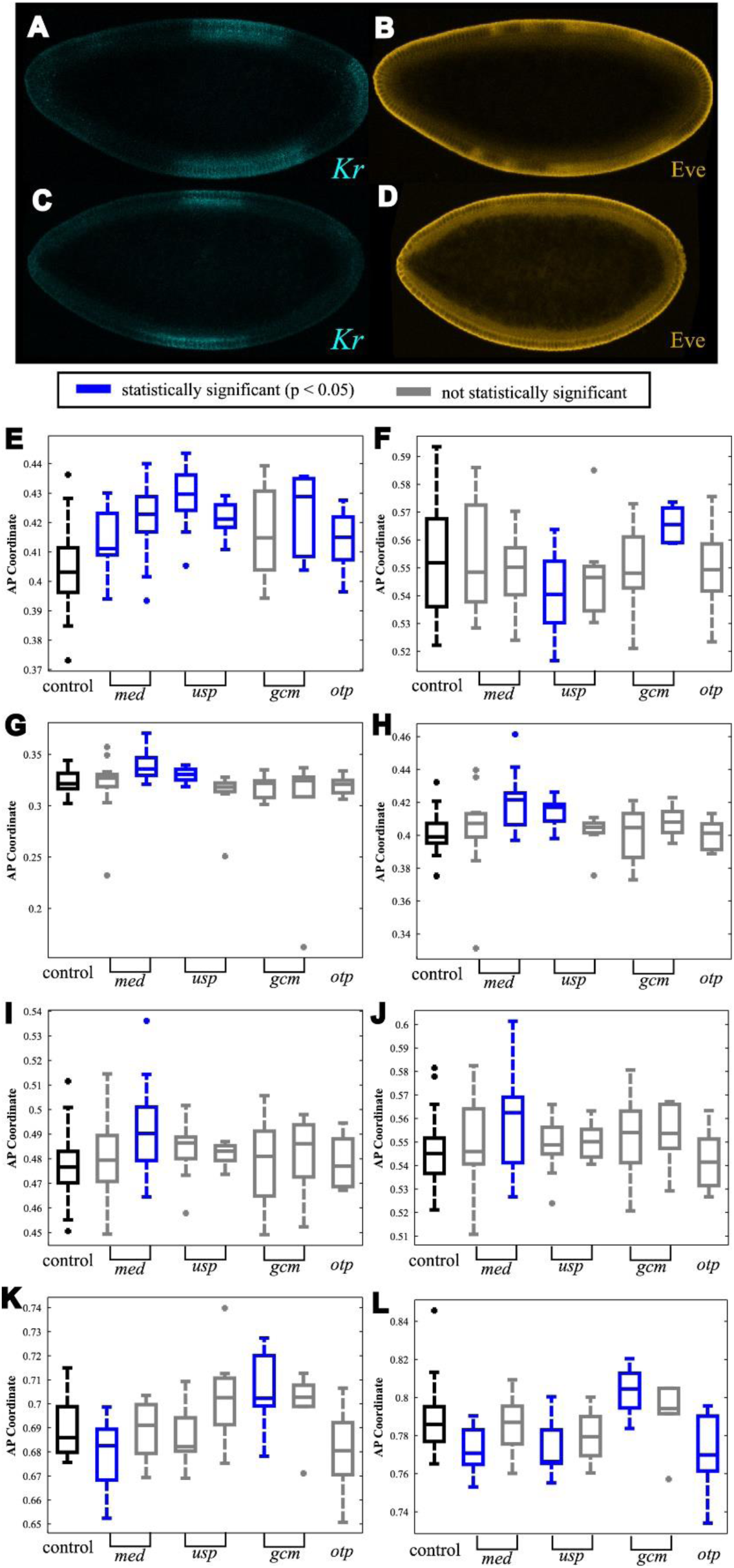
Shifts in *Kr* and Eve seen in mutants. In *pan* mutant BS26743 (A) *Kr* and (B) Eve expression. In mutant BS22312, (C) *Kr* and (D) Eve expression. Variation in positioning of (E) *Kr* anterior border, (F) *Kr* posterior border, (G) Eve stripe 1, (H) Eve stripe 2, (I) Eve stripe 3, (J) Eve stripe 4, (K) Eve stripe 6, and (L) Eve stripe 7.

### Shifts in Eve stripes and Kr borders are observed in usp mutants

The maternal gene *ultraspiracle* (*usp*) was found to effect expression of *Kr* and Eve in the early embryo. *usp* is expressed throughout the early embryo, however only weak expression remains by mid-NC14 [49–51]. Two fly lines mutated for *usp*, one amorphic and one hypomorphic, were found to produce shifts in the borders of *Kr* and Eve stripes 1, 2 and 7 (Fig. 4, S3 Table, S4 Table, and Materials and Methods). *usp* was tested because PWM analysis points to a binding site at the SNP in the EveC enhancer. This SNP was correlated with a shift in Eve stripe 5. Using the reporter constructs, this EveC enhancer produces *lacZ* expression in the anterior pole, which is lost when the SNP is mutated, and in stripes throughout the embryo where expression increases by mutating the SNP (Fig. 3). This expression due to the EveC enhancer (and the changes in expression when the SNP within the enhancer is mutated) are consistent with regions of the embryo where *Kr* and Eve stripe 1, 2, and 7 are located. While this effect of *usp* does not directly address why Eve stripe 5 was correlated with the SNP in its genomic context, perhaps further regulatory mechanisms effect the read-out of the SNP besides *usp*.

### *medea* results in subtle shifts in *Kr* and Eve throughout the AP axis

ChIP data (supported by PWM analysis) suggest a binding site for Medea (*Med*) in either EveA or EveB enhancer, at the SNP/indel contained within these reporter constructs. The SNP/indel within the EveA and EveB putative enhancers were correlated with a shift in Eve stripe 6 and stripe 5 respectively. Testing of *medea* mutants showed shifts in both the *Kr* anterior border and positioning of all Eve stripes (Fig. 4, S3 Table, S4 Table, and Materials and Methods). These effects spread throughout the entire embryo are consistent with expression observed due to the mutant EveA putative enhancer. Since *med* is maternally deposited and found ubiquitously throughout the early embryo these effects are consistent with the region where Medea is known to be present [52]. The main role of Medea is as an effector molecule for the Bone Morphogenetic Protein (BMP) pathway, which patterns the dorsal axis of the embryo [52]. However, Medea, and other elements of the Dpp pathway, also affect Wingless signaling [53], which has an AP patterning role at this stage [54]. In particular, Mad, which partners with Medea in BMP signal transduction [52], interacts with the Wingless effector protein Pangolin [53], which was also identified in our screen (see below).

### Positioning of *Kr* and Eve stripe 7 are shifted in *otp* mutants

PWM analysis suggested an Orthopedia (otp) binding site is present at the SNP in the KrA enhancer. This enhancer drives expression at the anterior pole, which is lost when the SNP is mutated (Fig 4). The SNP in the KrA enhancer was found to be correlated with a shift in the anterior border of *Kr*. An *otp* RNAi knockdown fly line has a shift in the expression of the anterior border of *Kr* and in Eve stripe 7 (Fig. 4, S3 Table, S4 Table, and Materials and Methods). *otp* is a Hox gene which is known to be active following gastrulation [49–51,55]. However, RNAseq data have found transcripts in this time period (2–4 hour old embryos) [56]. This suggests some low level of *otp* expression that affects *Kr* and possibly also Eve expression.

### *pangolin* mutations result in large shifts in *Kr* and Eve expression

The most significant effects were observed for fly lines mutated for *pangolin* (*pan*). *pan* expression is expressed ubiquitously throughout the early embryo and was tested based on PWM analysis of the EveB enhancer. The EveB enhancer generates expression at the anterior pole; however the mutated reporter construct generates expression throughout the embryo. Two different mutant fly lines, one expressing an RNAi knockdown (BS26743) and one with an insertion in the gene (BS22312) were tested. The *Kr* and Eve patterns produced by these fly lines were variable within these mutant populations. Significant differences compared to wild-type patterns were observed, examples for each fly line are seen in Fig. 4. For the insertion fly line (Fig. 4C,D), approximately half (7 out of 13) of the embryos tested exhibited this pattern. Over a quarter of the flies (4 out of 14) tested in the RNAi knockdown line (Fig 4A,B) showed this large expansion of the *Kr* domain and subsequent disruption in the Eve pattern. This suggests *pan* is necessary for the proper positioning of *Kr* and Eve.

## Discussion

Typical methods to study GRNs include labor-intensitve single-gene analysis [10,11,57] and genome-wide studies (e.g., [12,13,58,59]). In each of these cases, either the laboratory strain, or overtly-perturbed mutants are studied. Given that even the best-studied enhancers need further dissection before we attain a full understanding of cis-regulation [16], we used an alternative approach that focused on using the wild-caught DGRP lines to uncover the causes of subtle variation. In both engineering and systems biology contexts, subtle differences may point to compensatory regulation [60–64]. Therefore, we were not concerned with characterizing subtle differences *per sé*, but instead, we leveraged them to discover novel actors and regulation. Such regulation is difficult to discern in the laboratory strain, as it operates invisibly until a disturbance variable (e.g., small variations in humidity or nutrition) upsets the system [64]. As such, it may be a mechanistic example of the notions of cryptic variation and buffering [65–69].

Previous work has taken advantage of natural variation among the DGRP fly lines to discover new genes involved in the phenotypic and quantitative differences between these lines [29–31]. In doing so, new genes responsible for effects within diverse systems have been determined. Here we demonstrated the ability to detect subtle variations in *Kr* and Eve expression patterns, which led usto identify candidate genomic variants for futher testing. Using reporter constructs, we were able to validate these putative regulatory regions containing these genomic variants and identified certain SNP/indels which are able to produce allele-specifc activity and therefore likely are at transcription factor binding sites. Through this analysis we identified novel transcription factors (*usp, med, gcm*, and otp) that, when mutated, produce subtle variations in the position of *Kr* and Eve stripes. These subtle variations are consistent with the variations seen when the SNP/indels suspected of being their binding sites are mutated.

In this manner we also identified *pangolin*, which is able to produce large variations in *Kr* and Eve. The results of these analyses points to a greater network of genes involved in the anterior-posterior patterning system. In addition, this study demonstrates the ability of a SNP/indel to produce subtle, yet identifiable variations in gene expression. The methodology used in this study can be applied to further studies using the DGRP fly lines. A larger sample of lines (or all DGRP lines) can be tested, which would allow for SNPs to be explored throughout the genome (at significant distance from the gene of interest). This can identify trans-acting factors which were previously difficult to identify.

## Materials and Methods Embryo

### Staining and Image Collection

Embryos 2–4 hrs after egg lay, were fixed using formaldehyde per standard protocols. Subsequently, fluorescent *in situ* hybridization was used to image RNA and protein expression per published protocols (per [70] with proteinase K treatment omitted). RNA probes for *lacZ* (biotin conjugated) and *Kr* (flourescein conjugated) were used. Primary antibodies to biotin (goat, anti-biotin, 1:5000, gift from Immunoreagents), Eve (mouse anti-Eve, 1:10; Developmental Studies Hybridoma Bank), and flourescein (rabbit, anti-flourescein, 1:500; ThermoFisher Scientific). Secondary antibodies used were Alexa Flour 488 donkey anti-rabbit (ThermoFisher Scientific), Alexa Flour 546 donkey anti-goat (ThermoFisher Scientific), and AlexaFlour 546 donkey anti-mouse (ThermoFisher Scientific). Images were taken using a Zeiss Confocal microscope. Quantification of *Kr* and Eve was performed on images taken at the mid-saggittal plane and analyzed using Matlab, see [71].

### Plasmid Construction

The putative enhancer plasmids were cloned into the Evep:lacZ vector (gift from [33]) using EcoRI, BglII, or AscI. Enhancer regions were amplified from yw genomic DNA using primers listed in S1 Table. Mutations were introduced into the reporter constructs using mutagenesis PCR (primers listed in S2 Table). All PCR was carried out using Q5 Polymerase (New England BioLabs).

### Fly lines

*yw* was used as a laboratory control strain. Natural variation fly lines were provided by Trudy MacKay [29]. The lines denoted in this paper as 1–13, are RAL41, RAL57, RAL105, RAL306, RAL307, RAL315, RAL317, RAL360, RAL705, RAL761, RAL765, RAL799, and RAL801(in that order). Mutants for suspect transcription factors were obtained from Blooming Stock Center: *Medea* (BS9033 [Med^1^] and BS9006 [Med^5^] from ethyl methanesulfonate mutagenesis)*,pangolin* (RNAi knockdown line BS26743 and knockdown by transposable insertion within gene BS22312), *ultraspiracle* (hypomorphic line BS4660 and amorphic line BS31414), *glial cells missing* (RNAi knockdown lines BS28913, BS31518, and BS31519), and *orthopedia* (RNAi knockdown line BS57582). The reporter construct and mutant reporter construct fly lines were created by injection and incorporation of plasmid constructs into the 68A4 landing site (yw; attP2 flies) for the KrA:lacZ, EveL:lacZ, EveI:lacZ, and EveK:lacZ injections were performed by Model System injections. KrB:lacZ, KrC:lacZ, KrD:lacZ,, EveA:lacZ, EveB:lacZ, EveC:lacZ, EveD:lacZ:, EveE:lacZ, EveF:lacZ, EveG:lacZ, EveH:lacZ, EveJ:lacZ injections were performed by Genetic Services, Inc. KrBmut:lacZ, EveBmut:lacZ, EveCmut:lacZ, Evep:lacZ and EveGmut:lacZ injections were performed by GenetiVision Inc.

### Identification of Novel Transcription Factors

Identification of novel transcription factors using position weight matrices was carried out for the region surrounding the SNP within the enhancers that generated expression *in vivo*. The position weight matrices used were obtained from [35–40,40–42]. Probability of a transcription factor binding site being present within a given series was calculated by multiplying the probability of the given nucleotide at each position in the sequence and dividing this by the probability in a random sequence (calculated from a 10,000 bp *Drosophila melanogaster* exon region). Transcription factors where the precence of a site at the SNP/indel has p < 0.0005 were explored. Chromatin immunoprecipitation data were taken from MacArthur et al, 2009 [35].

## Acknowledgements

We would like to thank Dr. MacKay for the fly lines used in this study and helpful discussions.

## Supporting information

**Figure S1:**
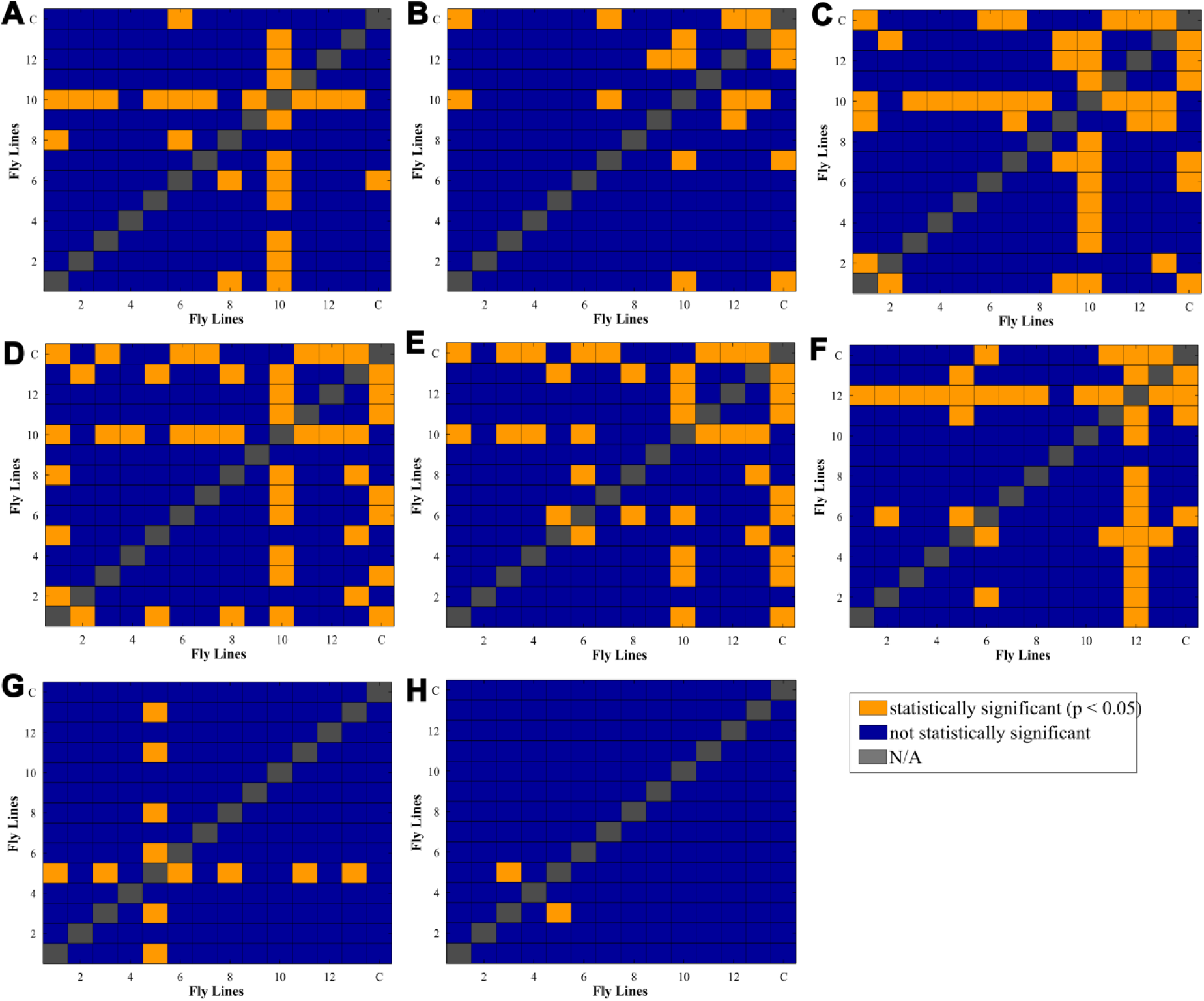
Pair-wise comparison of position of *Kr* and Eve in each of the fly lines. Results of post-hoc Tukey-Kramer test, where orange denotes statistically significant (p < 0.05) differences between the lines and blue denotes no statistically significant difference between the lines. The fly lines are in the order of: RAL150, RAL306, RAL307, RAL315, RAL317, RAL360, RAL41, RAL57, RAL705, RAL761, RAL765, RAL799, RAL801, and laboratory control; for (A) Kr Posterior, (B) Eve stripe 1, (C) Eve stripe 2, (D) Eve stripe 3, (E) Eve stripe 4, (F) Eve stripe 5, (G) Eve stripe 6, and (H) Eve stripe 7.

**Figure S2:**
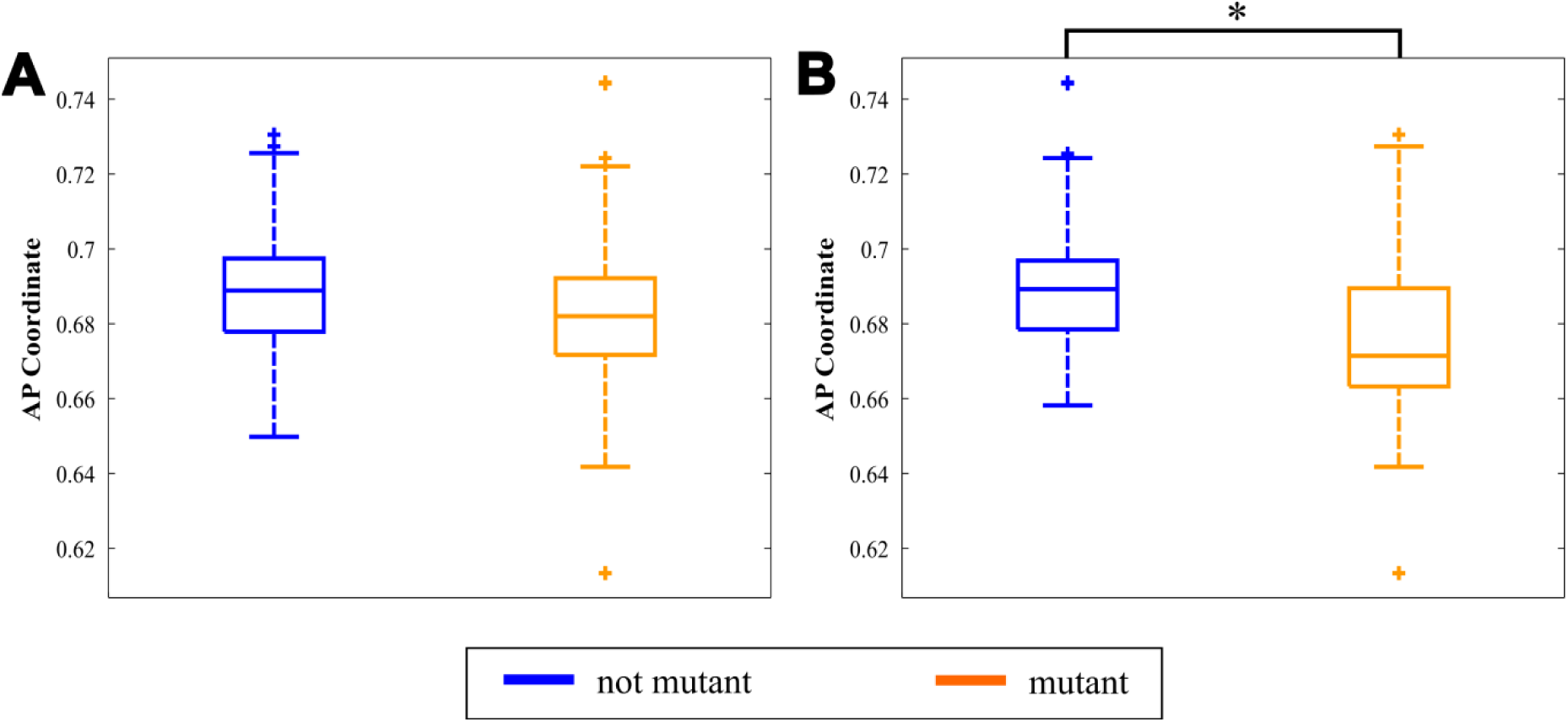
Example of Association Mapping. (A) Comparison of position of Eve stripe 6 in nonmutant lines (n_lines_ = 8, n_embryos_ = 88), compared to mutant lines (n_lines_ = 5, n_embryos_ = 88), for a nonsignificant SNP. (B) For a significant SNP (p = 0.045), this SNP shows a correlation (anterior shift in Eve stripe 6) between non-mutant lines (n_lines_ = 8, n_embryos_ = 123) and mutant lines (n_lines_ = 5, n_embryos_ = 53).

**Figure S3:**
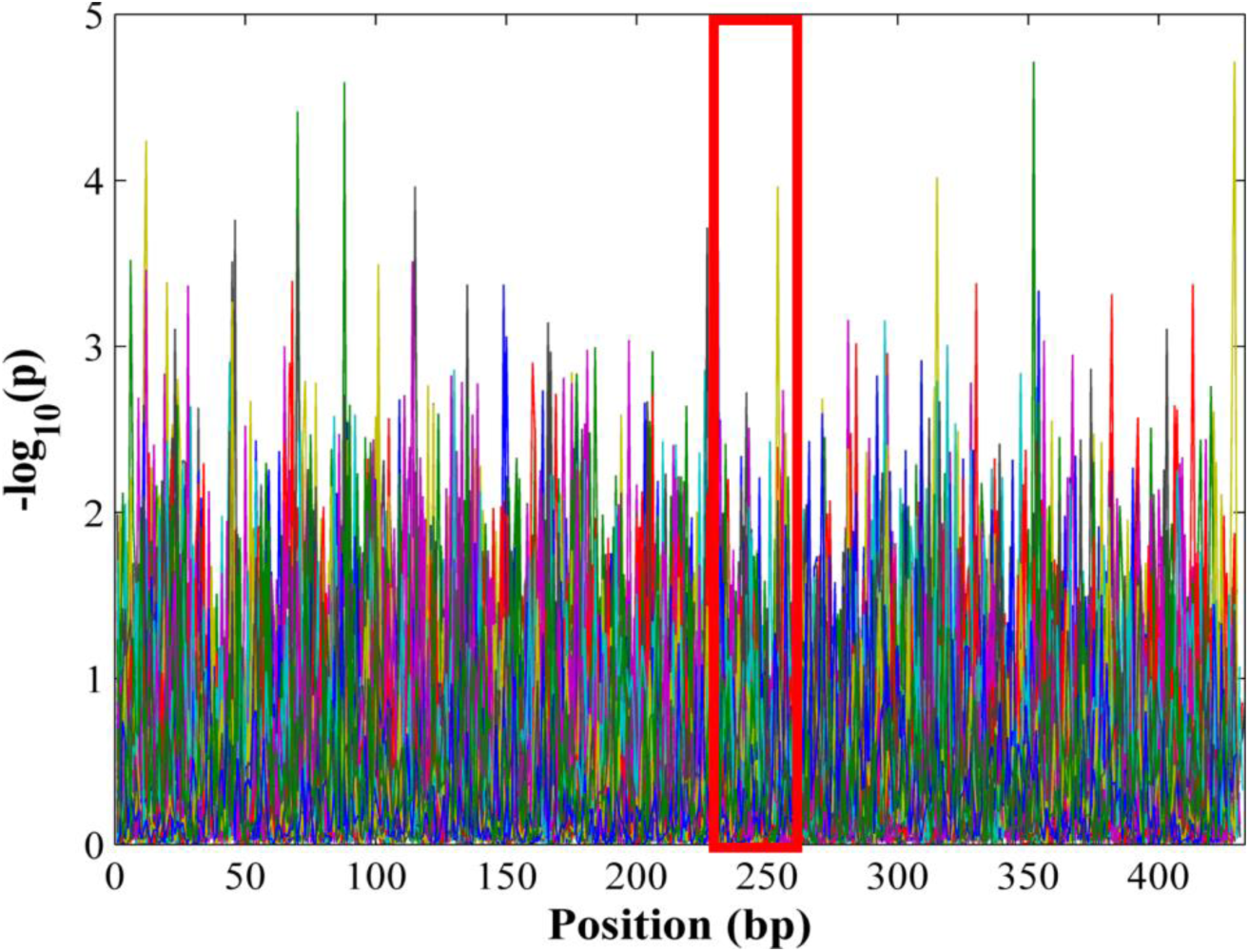
Position weight matrix analysis to find probable transcription factor binding sites. Analysis for EveA reporter construct, where each line denotes one transcription factor motif (only motifs shown to be present in one reporter construct are shown). The region directly surrounding the SNP is denoted by the red box.

**Figure S4:**
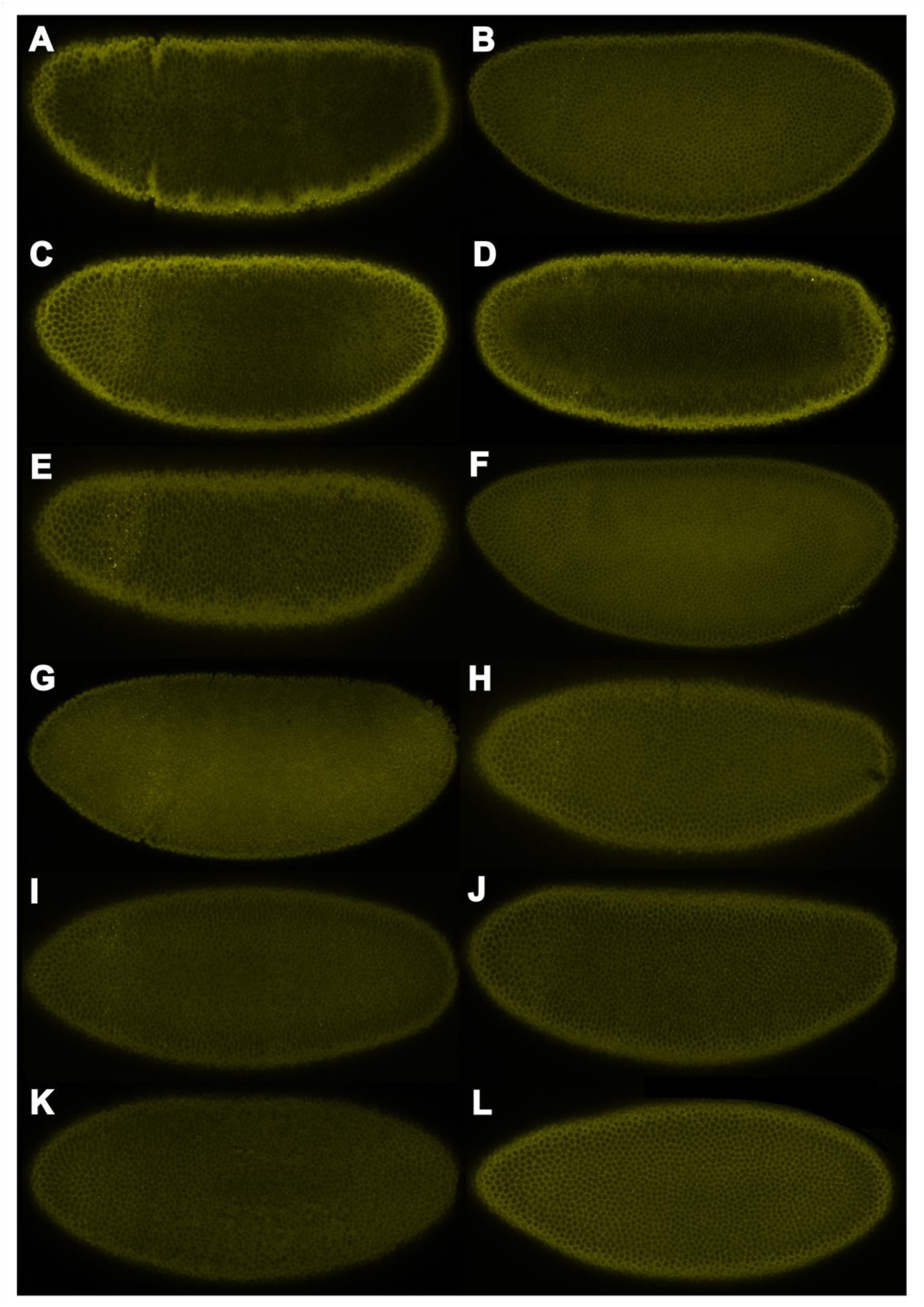
Expression of *lacZ* due to putative enhancer activity where no expression or expression due to only the minimal promoter is observed. (A) EveD, (B) EveE, (C) EveF, (D) EveG, (E) EveH, (F) EveI, (G) EveJ, (H) EveK, (I) EveL, (J) KrB, (K) KrC, and (L) KrD.

**Table S1:**
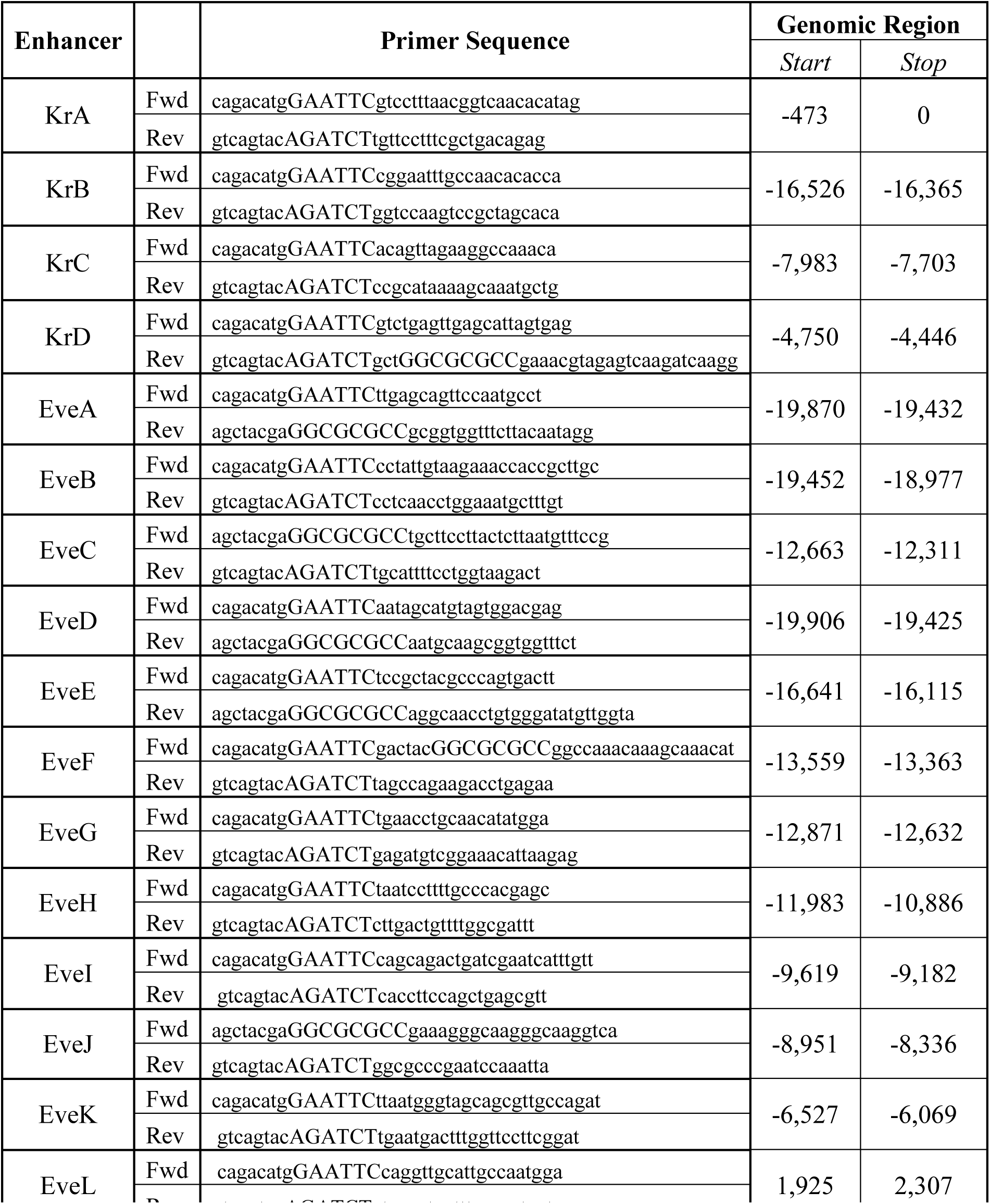
Primers used to amplify enhancers from genomic DNA. Restriction enzyme sites in capital letters. Genomic region of enhancer is shown compared to start of respective gene.

**Table S2:**
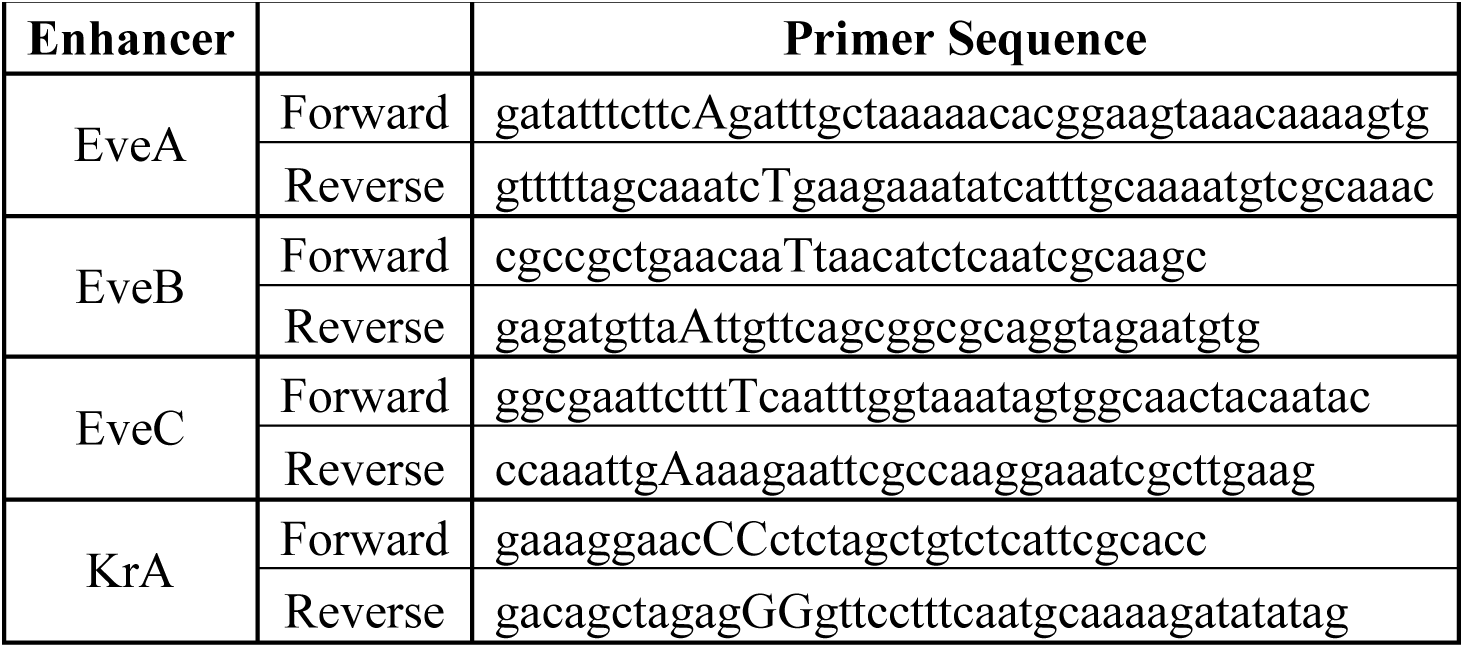
Sequence for Mutagenesis Primers. Mutation in capital letters.

**Table S3:**
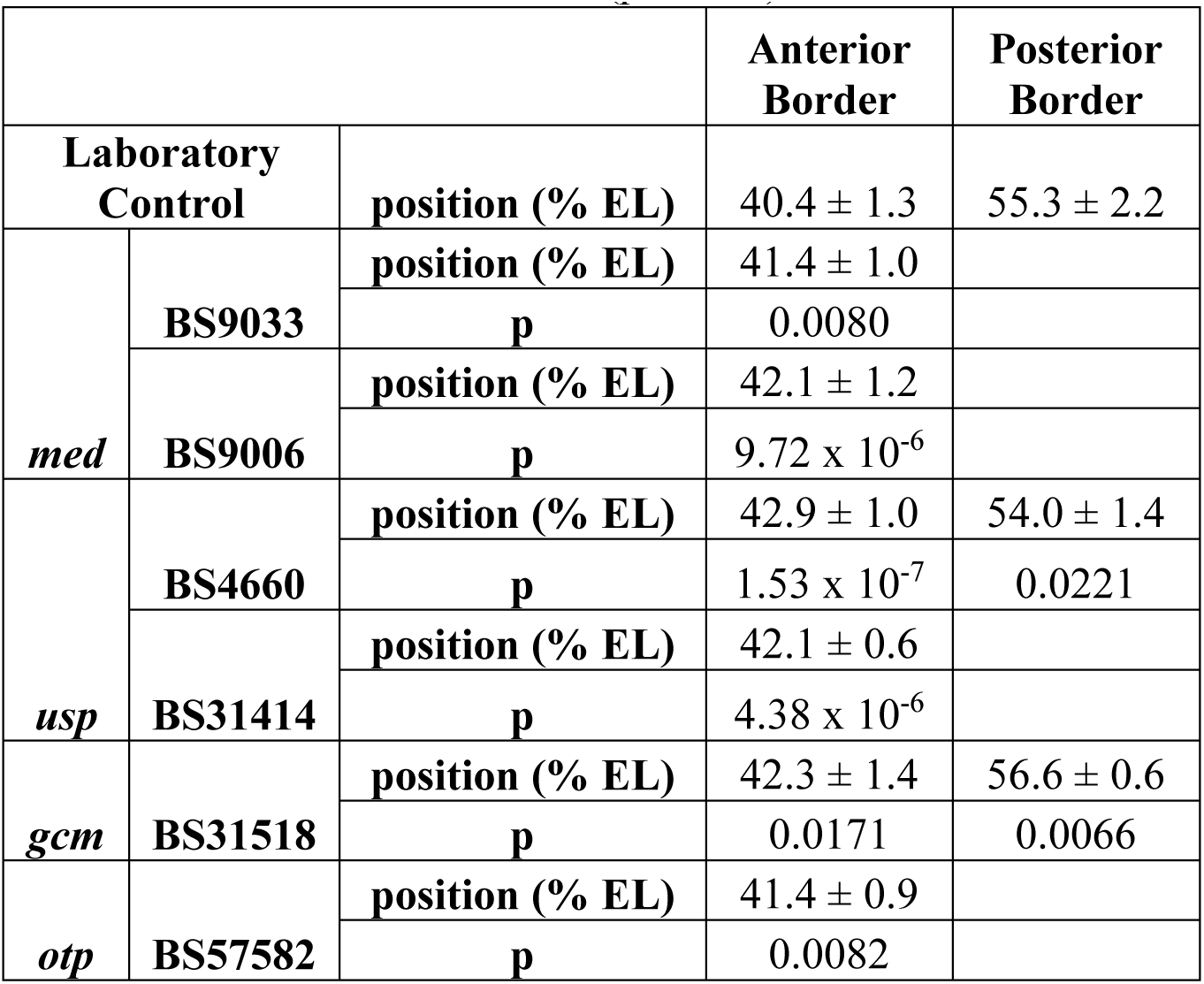
Shifts in positions of *Krüppel* in mutant fly lines. Shifts not statistical significant are not shown (p > 0.05).

**Table S4:**
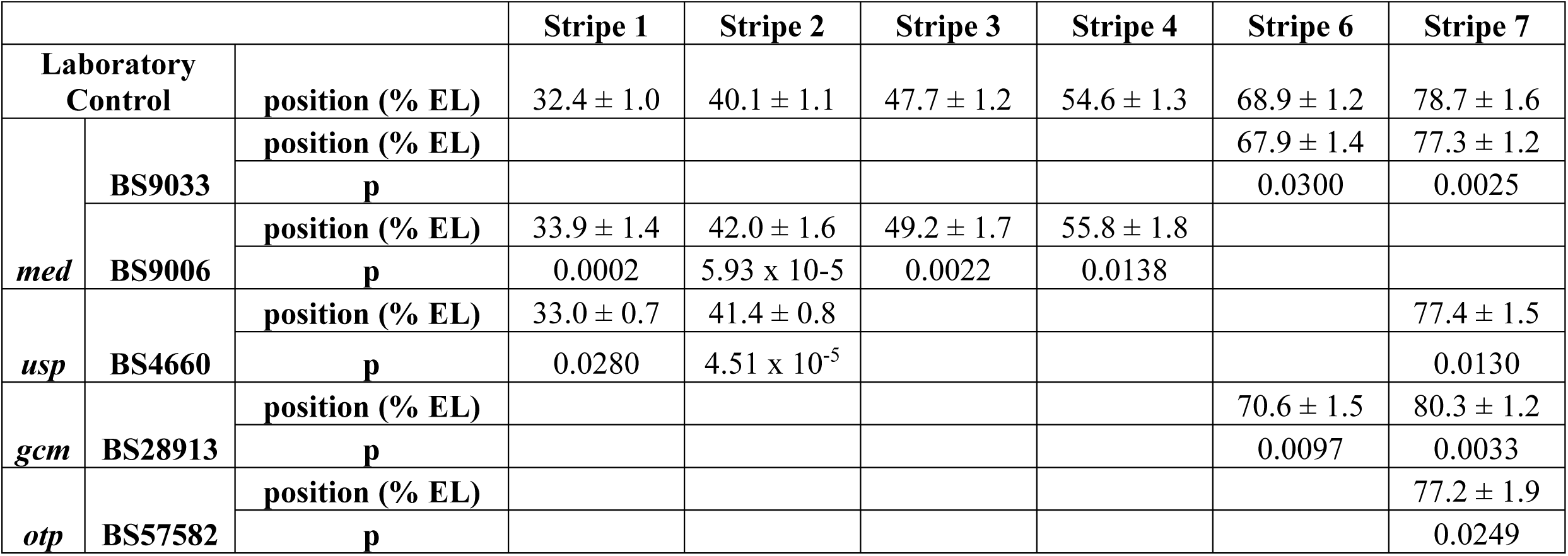
Shifts in the positions of Eve stripes in mutant fly lines. Shifts not statistical significant are not shown (p > 0.05).

|  |  |  | **Stripe 1** | **Stripe 2** | **Stripe 3** | **Stripe 4** | **Stripe 6** | **Stripe 7** |
| --- | --- | --- | --- | --- | --- | --- | --- | --- |
| **Laboratory Control** | | **position (% EL)** | $32.4 \pm 1.0$ | $40.1 \pm 1.1$ | $47.7 \pm 1.2$ | $54.6 \pm 1.3$ | $68.9 \pm 1.2$ | $78.7 \pm 1.6$ |
| *med* | **BS9033** | **position (% EL)** | | | | | $67.9 \pm 1.4$ | $77.3 \pm 1.2$ |
|  |  | **p** |  |  |  |  | 0.0300 | 0.0025 |
| | **BS9006** | **position (% EL)** | $33.9 \pm 1.4$ | $42.0 \pm 1.6$ | $49.2 \pm 1.7$ | $55.8 \pm 1.8$ | | |
| | | **p** | 0.0002 | $5.93 \times 10^{-5}$ | 0.0022 | 0.0138 | | |
| *usp* | **BS4660** | **position (% EL)** | $33.0 \pm 0.7$ | $41.4 \pm 0.8$ | | | | $77.4 \pm 1.5$ |
| | | **p** | 0.0280 | $4.51 \times 10^{-5}$ | | | | 0.0130 |
| *gcm* | **BS28913** | **position (% EL)** | | | | | $70.6 \pm 1.5$ | $80.3 \pm 1.2$ |
|  |  | **p** |  |  |  |  | 0.0097 | 0.0033 |
| *otp* | **BS57582** | **position (% EL)** | | | | | | $77.2 \pm 1.9$ |
|  |  | **p** |  |  |  |  |  | 0.0249 |

